# The 2D-structure of the *T. brucei* pre-edited RPS12 mRNA is not affected by macromolecular crowding

**DOI:** 10.1101/099200

**Authors:** W.-Matthias Leeder, Stephan Voskuhl, H. Ulrich Göringer

## Abstract

Mitochondrial transcript maturation in African trypanosomes requires RNA editing to convert nucleotide-deficient pre-mRNAs into translatable mRNAs. The different pre-mRNAs have been shown to adopt highly stable 2D-folds, however, it is not known whether these structures resemble the *in vivo* folds given the extreme “crowding” conditions within the mitochondrion. Here we analyze the effects of macromolecular crowding on the structure of the mitochondrial RPS12 pre-mRNA. We use polyethylene glycol as a macromolecular cosolute and monitor the structure of the RNA globally and with nucleotide resolution. We demonstrate that crowding has no impact on the 2D-fold and we conclude that the MFE-structure in dilute solvent conditions represents a good proxy for the folding of the pre-mRNA in its mitochondrial solvent context.

## Introduction

The folding of RNA molecules into compact, native structures or ensembles of structures is dictated by a set of first principle physicochemical forces.^1^ One of which is charge-compensation to overcome the electrostatic repulsion of the negatively charged phosphodiester backbone.^2–4^ Mono and divalent metal-ions at low to high millimolar concentrations contribute to this task^5^ and the effects of metal ion-radius and charge density have been studied in detail.^6–8^ Next to metal-ions, metabolites, polyamines and osmolytes have been shown to modulate RNA structure^9–13^ as well as high concentrations of macromolecules, which can occupy up to 30% of the total volume of a cellular compartment. This generates so-called “crowded” solvent conditions,^14–16^ which in general stabilize RNA 2D-and 3D-structures through the excluded volume effect and entropy perturbation of the folding landscape. ^5,17,18^

RNA editing in African trypanosomes describes a post-transcriptional modification reaction of mitochondrial pre-mRNAs that is characterized by the site-specific insertion and deletion of exclusively U-nucleotides (for a review see ref. 19). The reaction takes place within the single mitochondrion of trypanosomes, which represents the most crowded intracellular environment of eukaryotic cells. Intra-mitochondrial macromolecular concentrations reach up to 560g/L.^20,21^ Editing is catalyzed by a macromolecular protein complex, the 20S editosome, ^19^which interacts with 18 mitochondrial pre-mRNAs as substrates in the processing reaction. The different transcripts encode subunits of the mitochondrial electron transport and oxidative phosphorylation chains and have been characterized by several unusual features: First, the majority of pre-mRNAs lacks substantial sequence information (on average 45%), hence they require RNA editing to be converted into translatable mRNAs. Second, the different pre-mRNAs are typified by an extraordinarily high G-content (34%), which in two thirds of the cases are clustered in tracts of G-nucleotides (2≤G≤8). Third, ***in vitro*** chemical probing studies revealed that the different pre-mRNAs adopt extraordinarily stable 2D-structures approaching the stability of structural RNAs.^22,23^ Next to canonical base-pairing they contain pseudoknots and in many cases multiple G-quadruplex (GQ)-folds. However, it is not clear whether these 2D-structures resemble the ***in vivo*** folds given the extreme crowding conditions in the trypanosome mitochondrion. This is especially important since RNA editing ***in vitro*** has been shown to be sensitive to crowded solvent properties.^24^

Here we analyze the effect(s) of macromolecular crowding on the structure of the mitochondrial RPS12 pre-mRNA as an archetypical example of a mitochondrial transcript in African trypanosomes. We use high molecular mass polyethylene glycol (PEG) as a neutral macromolecular cosolute to mimic intra-mitochondrial solvent conditions and we monitor the structure of the RPS12 transcript by temperature-dependent UV-spectroscopy and by selective 2’-hydroxyl acylation analyzed by primer extension (SHAPE).

## Results and Discussion

To study the impact of macromolecular crowding on the structure of mitochondrial pre-mRNAs in *Trypanosoma brucei* we used the primary transcript of RPS12 as a representative model RNA. The pre-mRNA molecule is 325nt long. As a pan-edited transcript it is edited throughout its entire primary sequence with 132 U-nt inserted and 28 U’s deleted. The RNA has a G-nt-content of 27% and a purine/pyrimidine (R/Y)-ratio of 1.3. Its experimentally determined minimal free energy (MFE)-2D-structure calculates to a Gibbs free energy (ΔG) of −152kcal/mol with a base-paired *versus* single-stranded nucleotide ratio (r_bp/ss_) of 0.62. In addition, the RNA contains a pseudoknot.^22^ Since *in cell* structure probing experiments have successfully been performed only with abundant, cytosolic RNAs (see ref. 5,25-28) we decided to mimic the crowded, intra-mitochondrial solvent environment by using a chemically inert, synthetic cosolute such as polyethylene glycol (PEG).^29,30^ We used PEG with a mean molecular mass of 4000g/mol (PEG_4000_). The synthetic compound is characterized by a polymer crossover concentration (ϕ*) of 4% (w/w), which marks the transition from a semidilute to a crowded solvent regime.^31–33^ As a consequence, all experiments were performed at 6% (w/w) PEG_4000_.

As a first comparison, we measured UV-melting profiles of the RPS12 pre-mRNA in dilute and crowded solvent conditions. Representative normalized melting profiles and their 1^st^-derivatives (δA_260_/δT) are shown in Fig. 1A. At dilute buffer conditions (in the presence of 70mM Na^+^ and 2mM Mg^2+^), the pre-mRNA displays a complex melting profile with 5 distinct helix/coil transitions: Two dominant transitions with melting midpoints (T_m_-values) at 46°C and 78°C and three minor transitions at 30°C, 53°C and 62°C. Almost identical traces were recorded at crowded solvent conditions (Fig 1A). As demonstrated in the difference melting-curve in Fig. 1B, the two profiles superimpose perfectly at temperatures ≤40°C and deviate only slightly above 40°C. At that temperature the “crowded” profile shifts to higher temperature values, however only marginally with a ΔT of maximally 4°C (Fig. 1C). This indicates a very small structural stabilization of the transcript at volume-occupied solvent conditions. Since Mg^2^+-cations are known to drive the structural stabilization of RNAs we wondered whether any larger impact of the crowding agent was masked by the presence of Mg^2+^-cations. As a consequence, we re-analyzed the melting profile of RPS12-RNA in the absence of Mg^2+^. Representative normalized UV-melting curves and their 1^st^-derivatives (δA_260_/ δT) are shown in Fig. 1A. As expected, at dilute solvent conditions the melting profiles changed drastically: The T_m_ of the main transition shifted by 13°C from 44°C to 31°C and the formerly high temperature transition at 78°C disappeared altogether. However, as before, the PEG_4000_-induced stabilization was very small with a ΔT_m_ of 2°C for the main transition (Fig. 1A and Table 1A). This demonstrates that the crowding-driven stabilization of RPS12 RNA is by far weaker than the stabilization by divalent cations and that the impact on the overall structure of the transcript is minute.

**Figure 1.**
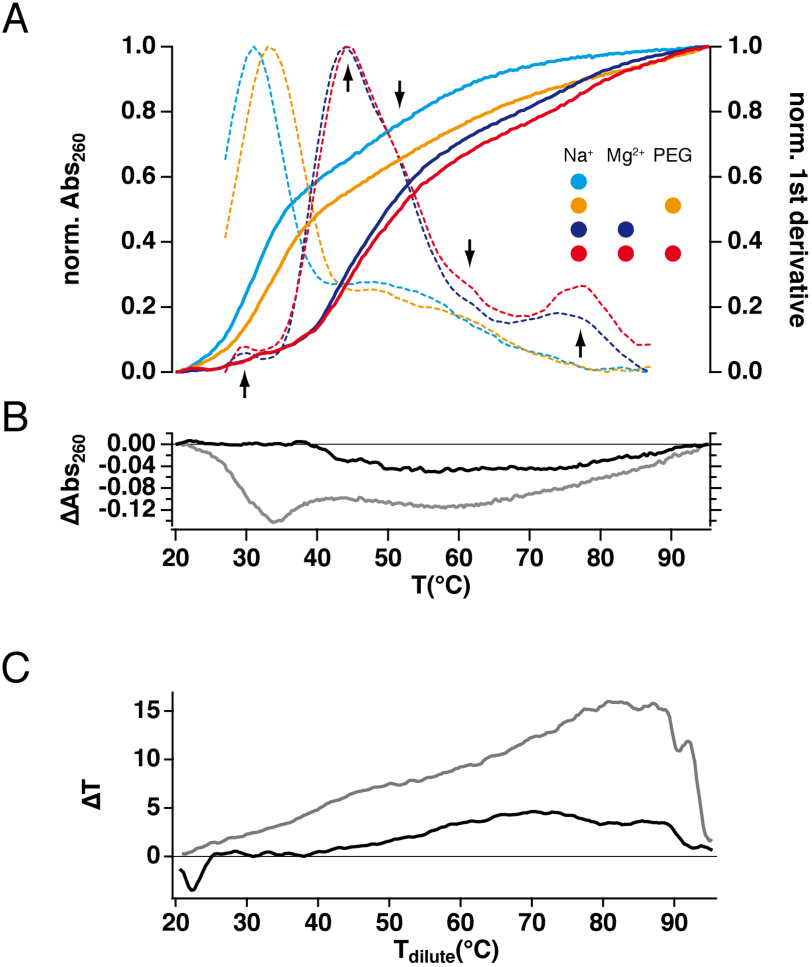
A: Normalized thermal UV-denaturation profiles of the RPS12 pre-mRNA from *Trypanosoma brucei* (colored, solid lines) at different buffered solvent conditions. Red/orange: crowded solvent conditions (6% (w/w) PEG_40_oo). Blue/cyan: dilute solvent conditions. Dark colors: 2mM Mg^2+^. Bright colors: no Mg^2^+. Dashed traces: Normalized 1^st^-derivatives (δA_260_/δT) of the different melting profiles. Five helix-coil transitions are marked with arrows B: Difference (Δ) UV-melting profiles (dilute minus crowded conditions) in the presence/absence of Mg^2^+-cations (black/grey). C: Shifting of the UV-denaturation profiles to higher temperatures expressed as ΔT *vs.* T of the dilute condition in presence/absence of Mg^2+^-cations (black/grey).

**Table 1.** A: Helix/coil transition temperatures (T_m_) of the *T. brucei* RPS12 pre-mRNA derived from the denaturation profiles shown in Fig. 1. Macromolecular crowding was mimicked by 6% (w/w) PEG_4000_. B: Summary of SHAPE-derived Gibb’s free energies (ΔG) and structural characteristics of the RPS12 transcript in dilute and crowded solvent conditions. In addition, mean and median SHAPE-reactivities are provided. C: Structural features of the RPS12 pre-mRNA in crowded and/or Mg^2+^-free solvent conditions compared to dilute solvent conditions at 2mM Mg^2+^. ΔΔG: difference Gibb’s free energies. sens: sensitivity. ppv: positive predictive value. For a comprehensive comparison see to Table S1.

|  |  |  |  |  |
| --- | --- | --- | --- | --- |
| **A** |  |  |  |  |
|  | dilute | crowded | dilute -Mg<sup>2+</sup> | crowded -Mg<sup>2+</sup> |
| $T_m$ (°C)<sub>main transition</sub> | 44.0 | 44.1 | 31.0 | 33.1 |
| **B** |  |  |  |  |
| $\Delta G_{37^\circ\text{C}}$ (kcal/mol) | -127.4 | -131.3 | -91.1 | -91.8 |
| fraction ss | 0.38 | 0.38 | 0.43 | 0.46 |
| fraction ds | 0.62 | 0.62 | 0.57 | 0.54 |
| mean reactivity | 0.45 | 0.39 | 0.57 | 0.61 |
| median reactivity | 0.35 | 0.32 | 0.35 | 0.37 |
| pseudoknotted | yes | yes | no | no |
| **C** |  |  |  |  |
|  | crowded vs.<br>dilute | dilute -Mg<sup>2+</sup> vs.<br>dilute | crowded -Mg<sup>2+</sup><br>vs. dilute |  |
| $\Delta T_m$ (°C) | <0.5°C | -13.0 | -11.0 | |
| $\Delta\Delta G_{37^\circ\text{C}}$ (kcal/mol) | -3.9 | 36.3 | 35.6 | |
| sens (%) | 100 | 56 | 52 |  |
| ppv (%) | 100 | 60 | 58 |  |

As a follow up of these experiments we analyzed the effects of macromolecular crowding by probing the structure of the RPS12 RNA with nucleotide resolution. For that we used selective 2’-hydroxyl acylation analyzed by primer extension (SHAPE).^34^ SHAPE monitors the local nucleotide flexibility of conformationally unrestrained nucleotides. *In toto* 282 nucleotides were interrogated using the four different solvent regimes depicted in Fig. 2. The nucleotide flexibility is measured in normalized SHAPE-units (SU) and as expected, the RPS12 transcript displays a complex reactivity pattern including highly reactive (>0.8SU) and almost unresponsive (<0.35SU) sequence regions in dilute solvent conditions.^22^ Sequence stretches with moderate (0.35≤SU≤0.8) to high flexibility are mostly clustered and alternate with unreactive sequence stretches (Fig. 2A). About half of the nucleotide positions are scored inflexible and roughly 10% are highly flexible (>0.8SU). The same probing experiments were performed at crowded conditions and the SHAPE-reactivity differences in the two solvent regimes were plotted as a difference (Δ_crowded-dilute_) SHAPE-profile (Fig. 2B). The two data sets are characterized by Pearson (*r*) and Spearman (*ρ*) correlation coefficients ≥0.92 (Table S1C/D) indicating that the structure of the RPS12 RNA is nearly identical in the two solvent regimes.

**Figure 2.**
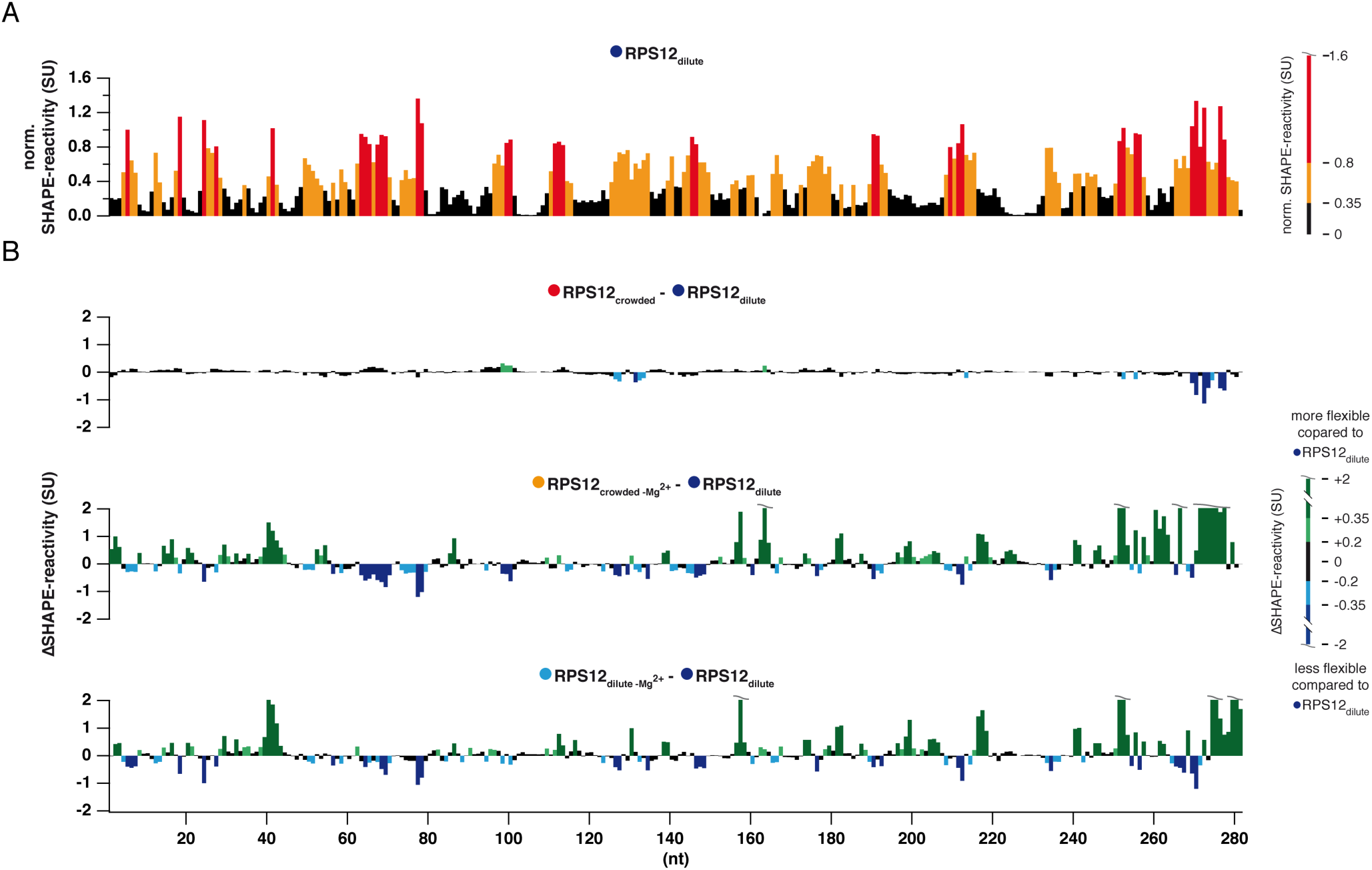
A: Normalized SHAPE-reactivity profile of the RPS12 pre-mRNA in dilute solvent conditions. Black: low (<0.35SU); orange: medium (0.35≤SU<0.8); red: high (≥0.8) normalized SHAPE-reactivities. SU: SHAPE-unit. nt: nucleotide(s). B: Difference (Δ) SHAPE-reactivity profiles of the RPS12 transcript at crowded and/or Mg^2+^-free solvent conditions. Green: nt-positions with increased SHAPE-reactivities (dark green: 0.35≤SU≤2; light green: 0.2≤SU<0.35). Blue: nt-positions with decreased SHAPE-reactivities (light blue: −0.2≥SU>−0.35; dark blue: −0.35≤SU≤−0.2. Black: non-responsive nt-positions (−0.2≤SU≥0.2).

As expected, a comparison of the SHAPE-profiles in the absence of Mg^2+^-ions resulted in a different picture. In the absence of the divalent cation both Δ SHAPE-profiles change throughout the entire primary sequence (Fig. 2C/D). About 20% of the nucleotides display reactivity-changes >|0.2|SU and an additional 33% show more than |0.35|SU difference. As a result, the data sets only correlate with correlation coefficients of *r*=0.27 (*ρ*=0.35) (crowded-Mg^2+^ vs. dilute) and *r*=0.4 (*ρ*=0.51) (dilute-Mg^2+^ vs. dilute) (Table S1C/D). This behavior translates to the 2D-structure level when the normalized SHAPE-reactivities are used as pseudo free energy constraints to guide the structure prediction. Fig. 3 shows the pseudoknotted 2D-structure of the RPS12 transcript in dilute and crowded solvent conditions. Molecular crowding has no effect on the transcript if Mg^2+^-ions are present. However, since the average and median nucleotide flexibility are slightly decreased at crowded solvent conditions this results in a decrease of the Gibbs free energy (ΔG) of −3.9kcal/mol (Tab. 1B/C). By contrast, in the absence of Mg^2+^-ions the two structures are characterized by a roughly 15% higher ΔG and the absence of the pseudoknot. Sixty percent of the nucleotides retain their structural context at all conditions studied (Fig. S2).

**Figure 3.**
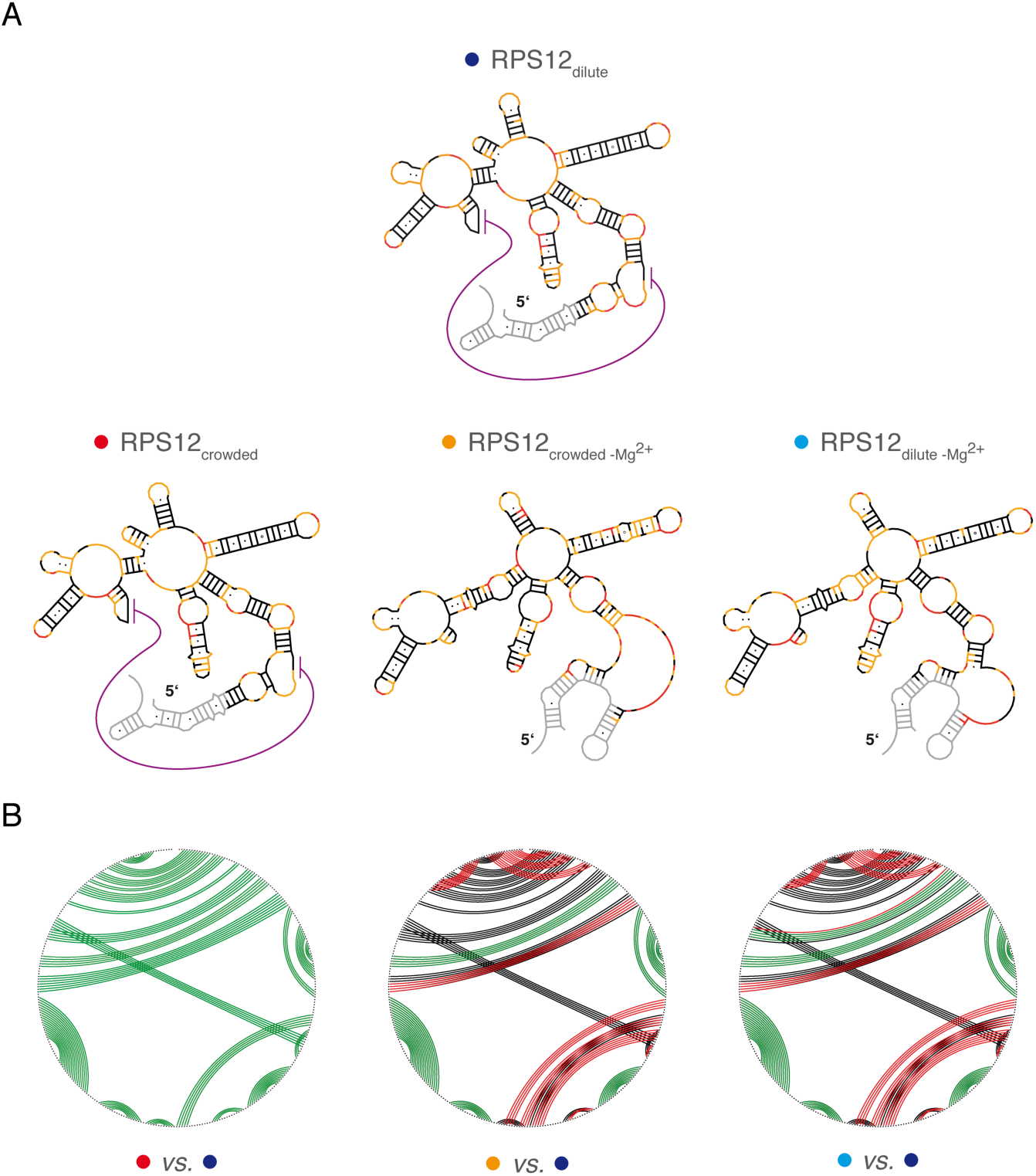
A: SHAPE-derived minimum free energy (MFE)-2D-structures of the RPS12 pre-mRNA - top: at dilute solvent conditions in the presence of 2mM Mg^2+^; bottom: at crowded solvent conditions in the presence/absence of 2mM Mg^2+^. B: Circle-plot comparison of SHAPE-derived 2D-structures of the RPS12 pre-mRNA. Base pairs (bp) are shown as colored lines. Green: bp present in both, dilute and crowded solvent conditions. Black: bp unique to dilute solvent conditions in the presence of Mg^2+^. Red: bp unique to the specific solvent conditions.

These results are in line with the published data of Soto *et al.*, 2007 and Tyrrell *et al.*, 2015 (see ref. 3,35). As expected, Mg^2+^-ions exert a stabilizing effect on the global fold of RNA molecules, which is reflected in a ΔT_m_ of 15°C and a ΔΔG of −36kcal/mol. The contributions of PEG-induced macromolecular crowding to the RNA-stability were less pronounced. Crowding conditions caused an increase of the main T_m_≤2°C and a decrease in the Gibb’s free energy of maximally 4kcal/mol. Similar trends have been described by Katari *et al.*, 2013 and by Kilburn *et al.*, 2013.^18,24^ The results obtained by SHAPE- and UV-thermal melting experiments in the absence of Mg^2+^-ions are at a first glance contradictory. The addition of PEG_4000_ resulted in an increase of the T_m_ by about 2°C but had no effect on the ΔG. Crowding conditions on average even increased the local nucleotide flexibility and had a destabilizing effect on some formerly stable structural elements (Table 1 and Fig. 3). This can be explained by the excluded volume effect lowering the degrees of freedom for conformational sampling ultimately trapping the RNA in misfolded states.^18,35^ When Mg^2+^ is present, charge compensation by the ion-sphere and/or chelated Mg^2+^-ions lead to an electrostatic collapse of the RNA thereby generating a highly compact folding state that is further stabilized by the excluded volume effect.^3,4,17,36–40^ In the presence and absence of Mg^2+^-ions the crowding-induced stabilization of the RPS12 pre-mRNA over the entire temperature range most likely results from a perturbed folding landscape due to a reduced conformationally freedom.^18^ Based on the data we conclude that there is no difference in the 2D-structure of RPS12 pre-mRNA in dilute and crowded solvent conditions at physiological Mg^2^+-ion concentrations. This is in line with a recent study of Tyrrell *et al.*, 2015.^35^ The authors used SHAPE to compare the folding of the aptamer domain of the adenine riboswitch at dilute, crowded and *in vivo* conditions in its free and ligand-bound state. They demonstrated that the 2D-structure of the ligand-free aptamer is recapitulated correctly at all conditions despite less pronounced 3D-interactions *in vitro*. Furthermore, ligand addition resulted in the formation of a more compact conformation involving several higher order 3D-interactions that were invariantly mapped in dilute and crowded solvent conditions and inside the cell. The RPS12 transcript displays non-varying local nucleotide flexibilities as well as identical melting profiles at dilute and crowded conditions given Mg^2+^ is present. This suggests a highly stable and compact structure that obviously is not affected by volume exclusion effects. Thus, we conclude that the *in vitro* experiments properly reproduce the *in vivo* situation, at least on the level of RNA secondary structure.

## Experimental Procedures

### DNA cloning and RNA synthesis

The mitochondrial gene encoding ribosomal protein S12 (RPS12) was PCR-amplified from *T. brucei* Lister 427 genomic DNA (see ref. 41) using the following DNA-oligonucleotide primers (KpnI and SacI restriction endonuclease recognition/cleavage sites are underlined): RPS12_forw. GG<u>GGTACC</u>CTAATACAC-TTTTGATAACAAACTAAAG; RPS12_rev. CC<u>GAGCTC-</u>CCTACCAAACATAAATGAACCTG. The PCR-amplicon was cloned into the KpnI and SacI endonuclease restriction sites of phagemid pBSSKII^-^ (Invitrogen) and transcripts were generated by run off *in vitro* transcription from linearized plasmid templates using T7-RNA polymerase. RNAs were purified from non-incorporated NTP’s by size exclusion chromatography, EtOH-precipitated and dissolved in 10mM Tris/HCl pH 7.5, 1mM EDTA (TE).

### UV-melting curves

RPS12-transcripts were dissolved in 0.5×TE pH7.5 (50μL), heated to 95°C (2min) and snap cooled on ice before the addition of a concentrated folding buffer to yield a final volume of 0.5mL. Final buffer concentration were: 5mM Na-cacodylate pH6.8, 70mM NaCl and 2mM MgCl_2_ or 5mM Na-cacodylate pH6.8 and 30mM NaCl (−Mg^2+^). Volume occupied conditions were generated by supplementing the folding buffer with PEG_4000_ to yield a final concentration of 6% (w/w). This concentration exceeds the polymer crossover concentration ϕ^*^_PEG4000_=4% (w/w), which marks the transition from a semi-dilute to a crowded solvent regime. RNA concentrations were adjusted to A_260_=0.5. Denaturation/renaturation profiles were measured at 260nm between 20°C and 95°C at a heating/cooling rate of 1°C/min (data acquisition: 0.3 data points/°C). Melting temperatures (T_m_) were obtained from the maximum of the first derivatives (δA_260_/δT) of the melting curves.

### SHAPE-modification

The modification reagent 1–methyl–7–nitroisatoic anhydride (1M7) was synthesized as described.^42^ RPS12 pre-mRNA (0.1μM) was denatured by heating to 95°C (2min) followed by snap cooling on ice. RNA refolding was achieved by equilibration in 20mM HEPES pH7.5, 30mM KCl, 10mM MgCl_2_ (dilute solvent conditions) or 20mM HEPES pH7.5 and 30mM KCl (-Mg^2+^) for 30min at 27°C, which represents the optimal growth temperature of insect-stage trypanosomes. Crowding conditions were induced by a PEG_4000_-concentration of 6% (w/w). RNA samples were split and treated either with 3.5mM 1M7 in DMSO or the same volume of neat DMSO. Modification reactions were quenched after 70sec by the addition of an equal volume of water. RNAs were recovered by EtOH precipitation and desalted by size exclusion chromatography.

### Reverse transcription and data processing

Equimolar amounts of fluorescently labelled DNA oligonucleotide primer T3 reverse: 6-FAM/JOE/TAMRA-AATTAACCCTCACTAAAGGGAAC were annealed to 1M7-modified or unmodified RNA samples in 0.25xTE pH7.5 by heating to 95°C (2min), cooling to 50°C (10min) and snap cooling on ice. Reverse transcription was performed in 50mM Tris/HCl pH8.3, 75mM NaCl, 3mM MgCl_2_, 2.5mM DTT, 0.25mM each dNTP and 0.75U/mL RiboLock RNase inhibitor (Invitrogen). The reaction was started by pre-warming the samples for 1.5min prior to the addition of 5U/mL SuperScript III reverse transcriptase (Invitrogen). RPS12 pre-mRNA was reverse transcribed for 20min at 40°C. Sequencing reactions were carried out using unmodified RNA, fluorescently labelled DNA oligonucleotide primer and ddCTP or ddGTP at a final concentration of 0.125mM each. Reverse transcription was stopped by snap cooling and the addition of 0.1 volume of 4M NaOH followed by heating to 95°C (5min). Samples were pooled, EtOH precipitated and redissolved in HiDi^®^ formamid (ABI/Life technologies) for capillary electrophoresis. Raw electrophoretic traces were analysed using SHAPEfinder^43^ utilizing the boxplot approach to determine the number of statistical outliers. Normalized SHAPE-reactivities were the result of averaging a minimum of 3 independent experiments.

### SHAPE-directed RNA folding

Normalized SHAPE-reactivities were used as pseudo Gibbs free energy (ΔG)-values to guide the folding of the RPS12 pre-mRNA. The minimum free energy (MFE) 2D-structure of the RPS12 transcript was generated using the ShapeKnots routine (see ref. 44) included in RNAstructure 5.6 see ref. 45) utilizing the default parameters m=1.8kcal/mol; b=0.6kcal/mol and p1=0.35kcal/mol; p2=0.65kcal/mol. Structure comparisons were performed using CircleCompare.^45^ MFE-structures were compared in terms of their “sensitivity” (sens) and their positive predictive values (ppv): Sens=fraction of bp in the reference structure also present in the non-reference structure; ppv= fraction of bp in the non-reference structure also occurring in the reference structure. RPS12 pre-mRNA probed in dilute buffer at physiological *i.e.* 10mM Mg^2+^-ion concentrations served as a reference state if not indicated otherwise.

## Disclosure of Potential Conflicts of Interests

No potential conflicts of interest were disclosed.

## Acknowledgements

This work was supported by the German Research Foundation (DFG-SFB902) and the Dr. Illing Foundation for Molecular Chemistry to H.U.G.

## Supplemental Materials

Supplemental material: Figure S1, Figure S2, Table S1

## REFERENCES

01. Doudna JA, Cech TR. The chemical repertoire of natural ribozymes. Nature 2002; 418:222–228; PMID:12110898; http://dx.doi.org/10.1038/418222a

02. Draper DE, Grilley D, Soto AM. Ions and RNA folding. Annu Rev Biophys Biomol Struct 2005; 34:221–243; PMID:15869389; http://dx.doi.org/10.1146/annurev.biophys.34.040204.144511

03. Soto AM, Misra V, Draper DE. Tertiary structure of an RNA pseudoknot is stabilized by “diffuse” Mg^2^+ ions. Biochemistry 2007; 46:2973–2983; PMID: 17315982; http://dx.doi.org/10.1021/bi0616753

04. Leipply D, Draper DE. Dependence of RNA tertiary structural stability on Mg^2^+ concentration: interpretation of the Hill equation and coefficient. Biochemistry 2010; 49:1843–1853; PMID: 20112919; http://dx.doi.org/10.1021/bi902036j

05. Tyrrell J, McGinnis JL, Weeks KM, Pielak GJ. The cellular environment stabilizes adenine riboswitch RNA structure. Biochemistry 2013; 52:8777–8785; PMID: 24215455; http://dx.doi.org/10.1021/bi401207q

06. Manning GS. The molecular theory of polyelectrolyte solutions with applications to the electrostatic properties of polynucleotides. Q Rev Biophys 1978; 11:179–246; PMID: 353876

07. Misra VK, Draper DE. Mg(2+) binding to tRNA revisited: the nonlinear Poisson-Boltzmann model. J Mol Biol 2000; 299:813–825; PMID: 10835286; http://dx.doi.org/10.1006/jmbi.2000.3769

08. Lambert D, Leipply D, Shiman R, Draper DE. The influence of monovalent cation size on the stability of RNA tertiary structures. J Mol Biol 2009; 390:791–804; PMID:19427322; http://dx.doi.org/10.1016/j.jmb.2009.04.083

09. Miyamoto S, Kashiwagi K, Ito K, Watanabe S, Igarashi K. Estimation of polyamine distribution and polyamine stimulation of protein synthesis in *Escherichia coli*. Arch Biochem Biophys 1993; 300:63–68; PMID: 7678729; http://dx.doi.org/10.1006/abbi.1993.1009

10. Lambert D, Draper DE. Effects of osmolytes on RNA secondary and tertiary structure stabilities and RNA-Mg^2^+ interactions. J Mol Biol 2007; 370:993–1005; PMID:17555763; http://dx.doi.org/10.1016/j.jmb.2007.03.080

11. Bennett BD, Kimball EH, Gao M, Osterhout R, Van Dien SJ, Rabinowitz JD. Absolute metabolite concentrations and implied enzyme active site occupancy in *Escherichia coli*. Nat Chem Biol 2009; 5:593-599; PMID:19561621; http://dx.doi.org/10.1038/nchembio.186

12. Lambert D, Leipply D, Draper DE. The osmolyte TMAO stabilizes native RNA tertiary structures in the absence of Mg^2^+: evidence for a large barrier to folding from phosphate dehydration. J Mol Biol 2010; 404:138–157; PMID:20875423; http://dx.doi.org/10.1016/j.jmb.2010.09.043

13. Trachman RJ III, Draper DE. Comparison of diamine and Mg^2^+ interactions with RNA tertiary structures: similar vs. differential effects on the stabilities of diverse RNA folds. Biochemistry 2013; 52:5911–5919; PMID: 23899366; http://dx.doi.org/10.1021/bi400529q

14. Zimmerman SB, Trach SO. Estimation of macromolecule concentrations and excluded volume effects for the cytoplasm of *Escherichia coli*. J Mol Biol 1991; 222:599–620; PMID: 1748995

15. Ellis RJ. Macromolecular crowding: obvious but underappreciated. Trends Biochem Sci 2001; 26:597–604; PMID:11590012; http://dx.doi.org/10.1016/S0968-0004(01)01938-7

16. Thirumalai D, Klimov DK, Lorimer GH. Caging helps proteins fold. Proc Natl Acad Sci USA 2003; 100:11195–11197; PMID: 14506295; http://dx.doi.org/10.1073/pnas.2035072100

17. Kilburn D, Roh JH, Guo L, Briber RM, Woodson SA. Molecular crowding stabilizes folded RNA structure by the excluded volume effect. J Am Chem Soc 2010; 132:8690–8696; PMID:20521820; http://dx.doi.org/10.1021/ja101500g

18. Kilburn D, Roh JH, Behrouzi R, Briber RM, Woodson SA. Crowders perturb the entropy of RNA energy landscapes to favor folding. J Am Chem Soc 2013; 135:10055–10063; PMID:23773075; http://dx.doi.org/10.1021/ja4030098

19. Göringer HU. ’Gestalt,’ composition and function of the *Trypanosoma brucei* editosome. Annu Rev Microbiol 2012; 66:65–82; PMID:22994488; http://dx.doi.org/10.1146/annurev-micro-092611-150150

20. Srere PA. The infrastructure of the mitochondrial matrix. Trends Biochem Sci 1980; 5:120–121; PMID:6409145

21. Harve KS, Lareu R, Rajagopalan R, Raghunath M. Understanding how the crowded interior of cells stabilizes DNA/DNA and DNA/RNA hybrids-in silico predictions and *in vitro* evidence. Nucleic Acids Res 2010; 38:172–181; PMID:19854935; http://dx.doi.org/10.1093/nar/gkp884

22. Leeder WM, Voigt C, Brecht M, Göringer HU. The RNA chaperone activity of the *Trypanosoma brucei* editosome raises the dynamic of bound pre-mRNAs. Sci Rep 2016; 6:19309; PMID: 26782631; http://dx.doi.org/10.1038/srep19309

23. Leeder WM, Hummel NF, Göringer HU. Multiple G-quartet structures in pre-edited mRNAs suggest evolutionary driving force for RNA editing in trypanosomes. Sci Rep 2016; 6:29810; PMID: 27436151; http://dx.doi.org/10.1038/srep29810

24. Katari VS, van Esdonk L, Göringer HU. Molecular crowding inhibits U-insertion/deletion RNA editing *in vitro:* consequences for the *in vivo* reaction. PLoS One 2013; 8:e83796; PMID:24376749; http://dx.doi.org/10.1371/journal.pone.0083796

25. Spitale RC, Crisalli P, Flynn RA, Torre EA, Kool ET, Chang HY. RNA SHAPE analysis in living cells. Nat Chem Biol 2013; 9:18–20; PMID: 23178934; http://dx.doi.org/10.1038/nchembio.1131

26. Ding Y, Tang Y, Kwok CK, Zhang Y, Bevilacqua PC, Assmann SM. *In vivo* genome-wide profiling of RNA secondary structure reveals novel regulatory features. Nature 2014; 505:696–700; PMID:24270811; http://dx.doi.org/10.1038/nature12756

27. Rouskin S, Zubradt M, Washietl S, Kellis M, Weissman JS. Genome-wide probing of RNA structure reveals active unfolding of mRNA structures *in vivo*. Nature 2014; 505:701–705; PMID: 24336214; http://dx.doi.org/10.1038/nature12894

28. McGinnis JL, Liu Q, Lavender CA, Devaraj A, McClory SP, Fredrick K, Weeks KM. In-cell SHAPE reveals that free 30S ribosome subunits are in the inactive state. Proc Natl Acad Sci USA 2015; 112:2425–2430; PMID: 21776997; http://dx.doi.org/10.1073/pnas.1411514112

29. Minton AP. The influence of macromolecular crowding and macromolecular confinement on biochemical reactions in physiological media. J Biol Chem 2001; 276:10577–10580; PMID: 11279227; http://dx.doi.org/10.1074/jbc.R100005200

30. Chebotareva NA, Kurganov BI, Livanova NB. Biochemical effects of molecular crowding. Biochem (Moscow) 2004; 69:1239–1251; PMID:15627378; http://dx.doi.org/10.1007/PL00021763

31. De Gennes P. Scaling concepts in polymer physics. Ithaca (NY): Cornell University Press; 1979

32. Kozer N, Schreiber G. Effect of crowding on protein-protein association rates: Fundamental differences between low and high mass crowding agents. J Mol Biol 2004; 336:763–774; PMID:15095986; http://dx.doi.org/10.1016/j.jmb.2003.12.008

33. Kozer N, Kuttner YY, Haran G, Schreiber G. Protein-Protein association in polymer solutions: from dilute to semidilute to concentrated. Biophys J 2007; 92:2139–2149; PMID:17189316; http://dx.doi.org/10.1529/biophysj.106.097717

34. Merino EJ, Wilkinson KA, Coughlan JL, Weeks KM. RNA structure analysis at single nucleotide resolution by selective 2’-hydroxyl acylation and primer extension (SHAPE). J Am Chem Soc 2005; 127:4223–4231; PMID: 15783204; http://dx.doi.org/10.1021/ja043822v

35. Tyrrell J, Weeks KM, Pielak GJ. Challenge of mimicking the influences of the cellular environment on RNA structure by PEG-induced macromolecular crowding. Biochemistry 2015; 54:6447–6453; PMID: 26430778; http://dx.doi.org/10.1021/acs.biochem.5b00767

36. Heilman-Miller SL, Pan J, Thirumalai D, Woodson SA. Role of counterion condensation in folding of the *Tetrahymena* ribozyme. II. Counterion-dependence of folding kinetics. J Mol Biol 2001a; 309:57–68; PMID:11491301; http://dx.doi.org/10.1006/jmbi.2001.4660

37. Heilman-Miller SL, Thirumalai D, Woodson SA. Role of counterion condensation in folding of the *Tetrahymena* ribozyme. I. Equilibrium stabilization by cations. J Mol Biol 2001b; 306:1157–1166; PMID:11237624; http://dx.doi.org/10.1006/jmbi.2001.4437

38. Thirumalai D, Lee N, Woodson SA, Klimov D. Early events in RNA folding. Annu Rev Phys Chem 2001; 52:751–762; PMID: 11326079; http://dx.doi.org/10.1146/annurev.physchem.52.1.751

39. Grilley D, Soto AM, Draper DE. Mg^2^+-RNA interaction free energies and their relationship to the folding of RNA tertiary structures. Proc Natl Acad Sci USA 2006; 103:14003–14008; PMID:16966612; http://dx.doi.org/10.1073/pnas.0606409103

40. Leipply D, Draper DE. Evidence for a thermodynamically distinct Mg^2^+ ion associated with formation of an RNA tertiary structure. J Am Chem Soc 2011; 133:13397–13405; PMID: 21776997; http://dx.doi.org/10.1021/ja2020923

41. Cross GA. Identification, purification and properties of clone-specific glycoprotein antigens constituting the surface coat of *Trypanosoma brucei*. Parasitology 1975; 71:393–417; PMID:645; http://dx.doi.org/10.1017/S003118200004717X

42. Turner R, Shefer K, Ares M Jr. Safer one-pot synthesis of the ‘SHAPE’ reagent 1-methyl-7-nitroisatoic anhydride (1m7). RNA 2013; 19:1857–1863; PMID: 24141619; http://dx.doi.org/10.1261/rna.042374.113

43. Vasa SM, Guex N, Wilkinson KA, Weeks KM, Giddings MC. ShapeFinder: a software system for high-throughput quantitative analysis of nucleic acid reactivity information resolved by capillary electrophoresis. RNA 2008; 14:1979–90; PMID: 18772246; http://dx.doi.org/10.1261/rna.1166808

44. Hajdin CE, Bellaousov S, Huggins W, Leonard CW, Mathews DH, Weeks KM. Accurate SHAPE-directed RNA secondary structure modeling, including pseudoknots. Proc Natl Acad Sci USA 2013; 110:5498–5503; PMID: 23503844; http://dx.doi.org/10.1073/pnas.1219988110

45. Reuter JS, Mathews DH. RNAstructure: software for RNA secondary structure prediction and analysis. BMC Bioinformatics 2010; 11:129; PMID: 20230624; http://dx.doi.org/10.1186/1471-2105-11-129

